# The respiratory virome and exacerbations in patients with chronic obstructive pulmonary disease

**DOI:** 10.1101/509919

**Authors:** Anneloes L. van Rijn, Sander van Boheemen, Ellen C. Carbo, Nikos Pappas, Igor Sidorov, Hailiang Mei, Marianne Aanerud, Per Bakke, Eric C.J. Claas, Tomas M. Eagan, Pieter S. Hiemstra, Aloys C.M. Kroes, Jutte J.C. de Vries

## Abstract

**Introduction:** Exacerbations are major contributors to morbidity and mortality in patients with chronic obstructive pulmonary disease (COPD), and respiratory bacterial and viral infections are an important trigger for the occurrence of such exacerbations. However, using conventional diagnostic techniques, a causative agent is not always found. Metagenomic next-generation sequencing (mNGS) allows analysis of the complete virome, but has not yet been applied in COPD exacerbations.

**Objectives:** To study the respiratory virome in nasopharyngeal samples during COPD exacerbations using mNGS.

**Study design:** 88 nasopharyngeal swabs from 63 patients from the Bergen COPD Exacerbation Study (2006-2010) were analysed by mNGS and in-house qPCR for respiratory viruses. Both DNA and RNA were sequenced simultaneously using an Illumina library preparation protocol with in-house adaptations.

**Results:** By mNGS, 23/88 samples tested positive. Sensitivity and specificity were both 96% for diagnostic targets (23/24 and 1067/1120, respectively). Viral pathogens only detected by mNGS were herpes simplex virus type 1 and coronavirus OC43. A positive correlation was found between Cq value and mNGS viral species reads (p=0.008). Patients with viral pathogens had lower percentages of bacteriophages (p<0.000). No correlation was found between viral reads (species and genus level) and clinical markers.

**Conclusions:** The mNGS protocol used was highly sensitive and specific for semi-quantitative detection of respiratory viruses. Excellent negative predictive value implicates the power of mNGS to exclude any infectious cause in one test, with consequences for clinical decision making. Reduced abundance of bacteriophages in COPD patients with viral pathogens implicates skewing of the virome, and speculatively the bacterial population, during infection.

## Introduction

Chronic obstructive pulmonary disease (COPD) is characterized by exacerbations with high morbidity and mortality, with over 65 million patients worldwide^1^. A COPD exacerbation is an acute event leading to worsening of the respiratory symptoms and is associated with a deterioration of lung function^2^. Exacerbations are mainly associated with infections, of which a large part is caused by viruses (22-64%) ^3–6^. However, in part of the exacerbations an etiologic agent is not detected.

Current routine virus diagnostics is based on polymerase chain reactions (PCR) and inherently the number of detectable pathogens is restricted to the ones included in the assay. Rare, mutated and pathogens with an uncommon clinical presentation will be missed, along with any new and currently unknown ones. Over the last decades, several previously unidentified viruses have been discovered as respiratory pathogen, including metapneumovirus^7^, middle-east respiratory syndrome coronavirus^8^ and human bocavirus^9^.

Metagenomic next generation sequencing (mNGS) is an innovative method which enables the detection of all genomes in a given sample. Proof of principle studies have shown that mNGS on respiratory samples can confirm and extend PCR results and deliver typing and resistance data at the same time^10–14^. The performance of mNGS in the clinical diagnostic setting, especially the positive and negative predictive value, has not yet been elucidated and is likely to differ per clinical syndrome and sample.

Previous data from reports on 16S rRNA analysis from the respiratory tract have led to increased insight in the microbiome in patients with COPD^15^. Changes in bacterial populations have been associated with exacerbation events and clinical phenotypes.^15^ However, these studies are intrinsically limited to analysis of the bacterial part of the microbiome.

So far only a few studies using shotgun metagenomics focus on the respiratory virome in children with acute respiratory infections^16,17^. In this study, we analyse the composition of the virome in adult patients with exacerbations of COPD.

## Objectives

The aim of this study was to correlate the respiratory virome in COPD patients as found by mNGS with qPCR and clinical data.

## Study design

### Patients

Patients with COPD were included in the Bergen COPD exacerbation study (BCES) between 2006 and 2010 in Bergen, Norway^18^. All patients lived in the Haukeland University Hospital district. Baseline data included amongst others exacerbation history, comorbidities, spirometry and Global Initiative for Chronic Obstructive Lung Disease (GOLD 2007) categorisation. Patients with an exacerbation were scheduled for an appointment with a study physician the next working day. During exacerbations, nasopharynx swabs were sampled and two different markers for the severity of the exacerbation were scored. After an exacerbation a control visit was scheduled. During the study period 154 patients had at least one exacerbation and in total 325 exacerbations were included in the Bergen COPD study, of which 88 exacerbation samples were tested in the current study.

### Sample selection

Nasopharyngeal samples were frozen and stored at −80°C. In total 88 nasopharyngeal samples of patients at the time of exacerbation were selected based on the availability of other samples (outside the current focus) and sent to the Leiden University Medical Center for further testing.

### Lab developed real-time PCR testing (qPCR)

The viral respiratory panel covered by the multiplex real-time PCR (qPCR) developed in our laboratory consists of coronavirus 229E, coronavirus HKU1, coronavirus NL63, coronavirus OC43, influenza A, influenza B, human metapneumovirus, parainfluenza 1-4 (differentiation with probes), respiratory syncytial virus, and rhinovirus^19^.

Total nucleic acids (NA) were extracted directly from 200 μl clinical sample, using the Total Nucleic Acid extraction kit on the MagnaPure LC system (Roche Diagnostics, Almere, the Netherlands) with 100 μL output eluate. Nucleic acid amplification and detection by real-time PCR was performed on a BioRad CFX96 thermocycler, using primers, probes and conditions as described previously^19^. Cq values were normalized using a fixed baseline fluorescence threshold.

### Metagenomic next generation sequencing (mNGS)

The metagenomics protocol used has been described previously^14^. In short, internal controls, Equine Arteritisvirus (EAV) for RNA and Phocid Herpesvirus-1 (PhHV) for DNA (kindly provided by prof. dr. H. G.M. Niesters, the Netherlands), were spiked in 200 μl of the virus transport medium in which the nasopharyngeal swab was stored. Nucleic acids were extracted directly from 200 μl clinical sample using the Magnapure 96 DNA and Viral NA Small volume extraction kit on the MagnaPure 96 system (Roche Diagnostics, Almere, The Netherlands) with 100 μL output eluate. For library preparation, 7 μl of nucleic acids were used, using the NEBNext^®^ Ultra™ Directional RNA Library Prep Kit for lllumina^®^, with several in-house adaptations to the manufacturers protocol in order to enable simultaneous detection of both DNA and RNA. The following steps were omitted: poly A mRNA capture isolation, rRNA depletion and DNase treatment step. This resulted in a single tube per sample throughout library preparation containing both DNA and RNA. Metagenomic sequencing was performed on an Illumina NextSeq 500 sequencing system (Illumina, San Diego, CA, USA), and approximately 10 million 150 bp paired-end reads per sample were obtained.

After quality pre-processing, sequencing reads were taxonomically classified with Centrifuge^20^ using an index constructed from NCBI’s RefSeq and taxonomy databases (accessed November 2017) with reference nucleotide sequences for the domains of viruses, bacteria, archaea, fungi, parasites, protozoa. Reads with multiple best matches were uniquely assigned to the lowest common ancestor (k=1 setting; previously validated^14^). Horizontal coverage (%) was determined using www.genomedetective.com^21^ version 1.111 (accessed 2018, December 30th).

### Assembly of the betacoronavirus

For samples with dubious or inconclusive classification results a *de novo* assembly was performed. Pre-processed short reads assigned to a higher taxonomic level of a suspected viral target were extracted and *de novo* assembled with SPAdes version 3.11.1^22^ into longer stretches of contiguous sequences (contigs). The resulting contigs were then run against the blast NCBI’s nucleotide (nt) database (accessed 2017) using blastn 2.7.1^23^. After identification of a putative target sequence, all the reads from the original sample were mapped against the identified best BLAST hit for further confirmation using BWA 0.7.17 software package^24^.

### Statistical analysis

Sensitivity, specificity, positive and negative predictive values were calculated based on 24 PCR positive and 1120 PCR negative target results of 88 samples.

Correlation between qPCR Cq value and logarithm of numbers of mNGS viral reads was tested with population Pearson correlation coefficient.

Potential correlations of mNGS data with clinical variables were tested as follows. Cq value/ viral reads and clinical parameters (exacerbation severity, duration of exacerbation or decrease/increase in Forced Expiratory Volume in 1 second (FEVi, control visit compared to baseline) were tested with one-way ANOVA and Kruskal-Wallis test when appropriate (depending on distribution). Comparison of the percentage of phages of all viral reads (after subtraction of the internal control EAV reads) between mNGS virus positive samples and negative samples was tested with Mann-Whitney U test, comparison with clinical parameters with Kruskal-Wallis test. Diversity of the virome in different patient groups was characterized by Shannon Diversity Index (H) and tested with Welch two sample t-test. Statistical analyses were performed using IBM SPSS Statistics version 23 software for Windows and R version 3.3.0. Differences at a p-value <0.05 were considered statistically significant.

### Ethical approval

The BCES study was approved by the ethical committee REK-Vest in Norway (REK number 165.08). The performance of additional testing, including mNGS, was approved by the medical ethics review committee of the Leiden University Medical Center (CME number B16.004).

## Results

### Patients and samples

In total 63 patients with 88 exacerbations were included with a median of one exacerbation per patient (range 1-5). Baseline patient characteristics and exacerbation characteristics are shown in tables 1 and 2 respectively.

**Table 1.** Baseline patient characteristics

|  | **Patients (n=63)** |
| --- | --- |
| Age median yrs (range) | 63.5 (46.6-74.5) |
| Male sex | 40 (64%) |
| BMI median (kg/m<sup>2</sup>) | 25 (15-39) |
| Body composition |  |
| Cachectic | 7 (11%) |
| Normal | 24 (38%) |
| Overweight | 22 (35%) |
| Obese | 10 (16) |
| Smoking |  |
| Never | 0 (0%) |
| Sometimes | 37 (59%) |
| Daily | 26 (41%) |
| GOLD stage |  |
| II (FEV<sub>1</sub> 50-80%) | 29 (46%) |
| III (FEV<sub>1</sub> 30-50%) | 27 (43%) |
| IV (FEV<sub>1</sub> <30%) | 7 (11%) |
| FEV<sub>1</sub> in % median (range) | 0.49 (0.23-0.74) |
| >1 exacerbation past 12 months | 16 (25%) |
| Inhalation steroids | 50 (79%) |

**Table 2.** COPD patient and exacerbation characteristics among patients having a viral or non-viral exacerbation.

|  | Virus*<br>detected<br>n=23 | Virus* not<br>detected<br>n=65 | p** |
| --- | --- | --- | --- |
| **Patient characteristics** |  |  |  |
| *Sex, %* |  |  | 0.21 |
| Women | 34.5 | 65.5 |  |
| Men | 22.0 | 78.0 |  |
| *smoking status, %* |  |  | 0.53 |
| Ex-smoker | 23.4 | 76.6 |  |
| Current-smoker | 29..3 | 70.7 |  |
| *GOLD stage (2007), %* |  |  | 0.35 |
| II (FEV<sub>1</sub> 50-80%) | 26.3 | 73.7 |  |
| III (FEV<sub>1</sub> 30-50%) | 30.8 | 69.2 |  |
| IV (FEV<sub>1</sub> < 30%) | 9.1 | 90.9 |  |
| *Frequent exacerbator, %* |  |  | 0.72 |
| No | 25.0 | 75.0 |  |
| Yes | 28.6 | 71.4 |  |
| *Using inhalation steroids, %* |  |  | 0.55 |
| No | 20.0 | 80.0 |  |
| Yes | 27.4 | 72.6 |  |
| *Age, mean yrs* | 63.7 | 64.9 | 0.10 |
| *BMI, mean kg/m<sup>2</sup>* | 27.0 | 25.9 | 0.92 |
| *FEV<sub>1</sub> in % predicted* | 49.3 | 47.5 | 0.48 |
| **Exacerbation characteristics** |  |  |  |
| *Exacerbation severity for entire exacerbation* |  |  | 0.75 |
| Mild (not requiring AB or oral steroids or hospitalization) | 14.3 | 85.7 |  |
| Moderate (requiring AB or oral steroids) | 26.9 | 73.1 |  |
| Severe (Emergency room or hospital admission) | 28.6 | 71.4 |  |
| *Self-reported exacerbation severity at time of study sampling* |  |  | 0.64 |
| Dyspnea unchanged or increased on errands outside home | 36.4 | 63.6 |  |
| Increased dyspnea doing housework | 26.5 | 73.5 |  |
| Increased dyspnea at rest | 28.6 | 71.4 |  |
| Must sit up at night due to dyspnea | 14.3 | 85.7 |  |
| *CRP (ng/mL) at time of study sampling<sup>†</sup>* | 32.5 | 34.2 | 0.27 |

\*qPCR target virus
\*\* Pearson chi-square for categorical variables and t-test for continuous variables
† missing data for 4 (1 virus positive, 3 virus negative) exacerbations

### Lab developed real-time PCR

Of the 88 samples, 23 (26%) tested positive with in-house PCR: 14 (61%) were rhinovirus positive, three influenza A, two coronavirus NL63, one coronavirus OC43, two parainfluenza 3 and one parainfluenza 4. Cq values ranged from 19-38 (Table 3).

**Table 3.** qPCR positive samples with respective mNGS results

| Samples | qPCR positive (%) | Cq values range | mNGS species positive (%) | mNGS species reads (range) | Coverage (% range) |
| --- | --- | --- | --- | --- | --- |
| All targets | 23/88 (26) | 19-38 | 23/88 (26) | 0-1,319,849 | 3-100 |
| Influenza A | 3/23 (13) | 29-36 | 3/23 (13) | 9-557 | 3-98 |
| Cov NL63 | 2/23 (9) | 32 | 2/23 (9) | 1,518-139,814 | 93-100 |
| Cov OC43 | 1/23 (4) | 27 | 2*/23 (4) | 1,319,849 | 99-99 |
| PIV3 | 2/23 (9) | 26-36 | 2/23 (9) | 58-275,644 | -** |
| PIV4 | 1/23 (4) | 24 | 1/23 (4) | 185,121 | 100-100 |
| Rhinovirus | 14/23 (61) | 19-38 | 13***/23 (57) | 0-40,409 |  |
|  |  |  | RV-A: 6/13 | 632-420,551 | 94-100 |
|  |  |  | RV-B: 2/13 | 38,421-409,480 | 100-100 |
|  |  |  | RV-C: 5/13 | 9,398-4,992,575 | 30-100 |

\*Retesting by qPCR confirmed the OC43 finding of mNGS
\*\* No coverage using GenomeDetective, Bowtie alignment confirmed Centrifuge mapping
\*\*\* Rhinovirus not detected with mNGS had PCR Cq value 38

### Metagenomic next generation sequencing

A median of 11 million (7,522,643-20,906,019) sequence reads per sample were obtained. Of the 11 million reads, approximately 94% were *Homo sapiens* reads, 2 % were bacterial and 0.1% viral (Table 4). No fungal reads were detected. A median of 2% of the reads could not be assigned to sequences in the Centrifuge index database (NCBI RefSeq).

**Table 4.** mNGS read counts

|  | Median | Min | Max |
| --- | --- | --- | --- |
| Root reads | 10,764,981 | 7,522,643 | 20,906,019 |
| % unassigned reads | 2 | 0.6 | 22 |
| Homo sapiens reads(% root) | 9,495,259 (94) | 2,510,133 | 18,672,027 |
| Bacterial reads (%root) | 233,472 (2) | 5,086 | 10,532,753 |
| Viral reads (% root) | 12,139 (0.1) | 500 | 1,544,795 |

### Comparison of mNGS to qPCR

Of the 23 qPCR positive samples, 22 tested positive with mNGS, resulting in a sensitivity of mNGS of 96%. Only one sample, that was rhinovirus positive by qPCR (Cq 38), could not be detected by mNGS (Table 3). Coverage of reference genomes was high (93-100%) with the exception of two samples: 30% coverage of rhinovirus C (1,401,120 mapped reads, 88,353-fold depth), and 3% coverage of influenza A virus (single genome segment, 8 mapped reads). Bowtie alignment confirmed the rhinovirus C mapping, but not the influenza A mapping. Additional viral pathogens detected by mNGS were herpes simplex virus type 1 (32,159 reads, 82% coverage, 36-fold depth) which is not in qPCR viral respiratory panel, in the sample with the 8 influenza virus reads, and a betacoronavirus. Of these 83,252 betacoronavirus reads, *de novo* assembly resulted in 3 contigs (size 30743, 274 and 232 bp respectively) with best BLAST hit coronavirus OC43 (reference genome accession AY391777). A coverage plot of all reads against this reference strain (Figure 1) showed good horizontal and vertical coverage (read coverage depth 428). The original OC43 qPCR amplification appeared to have been inhibited, and repeated OC43 qPCR confirmed the positive mNGS result (Cq 25).

**Figure 1.**
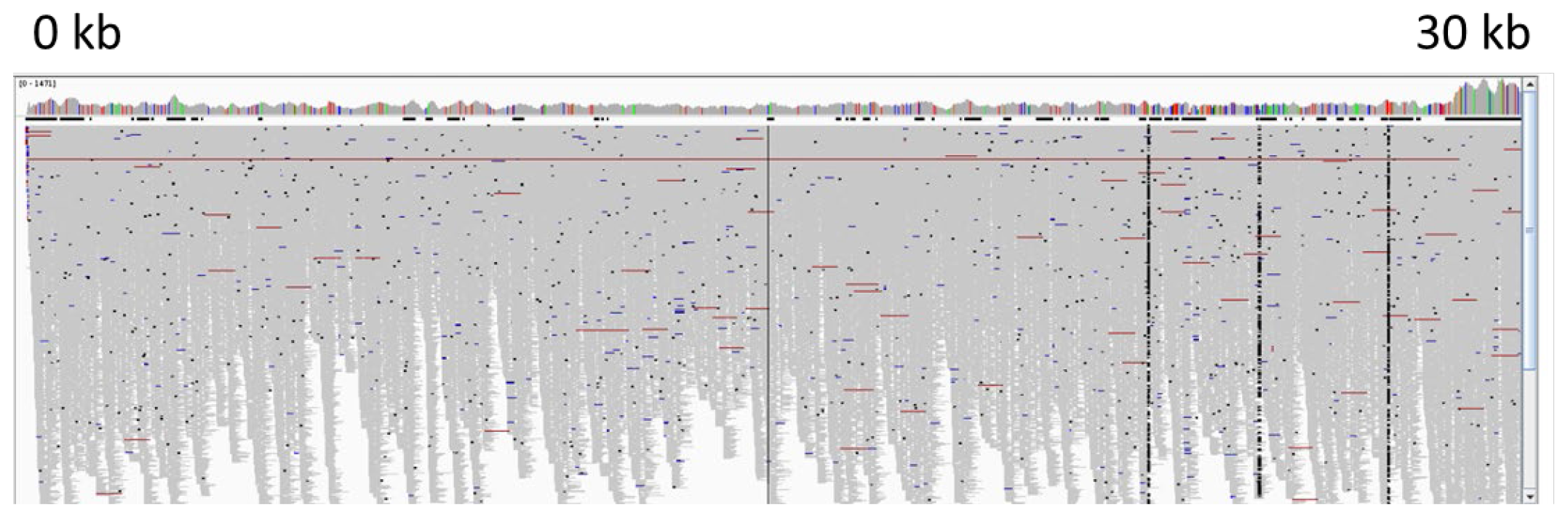
Coverage plot of betacoronavirus reads to coronavirus OC43 reference genome AY391777.1 (depth of coverage: 428).

### Sensitivity, specificity and predictive value

The sensitivity, specificity and predictive values of mNGS were calculated based on 24 PCR positive and 1120 PCR negative target results of 88 samples (Table 5). Calculations were made using different cut-off values of respectively >= 0, >15 and >50 mapped sequence reads. With a cut-off of >15 reads, the sensitivity was 92% and specificity 100%. With increasing cut-off levels, the positive predictive value (PPV) increased to 87%. The negative predictive value (NPV) was 100% for all cut-off levels.

**Table 5.**
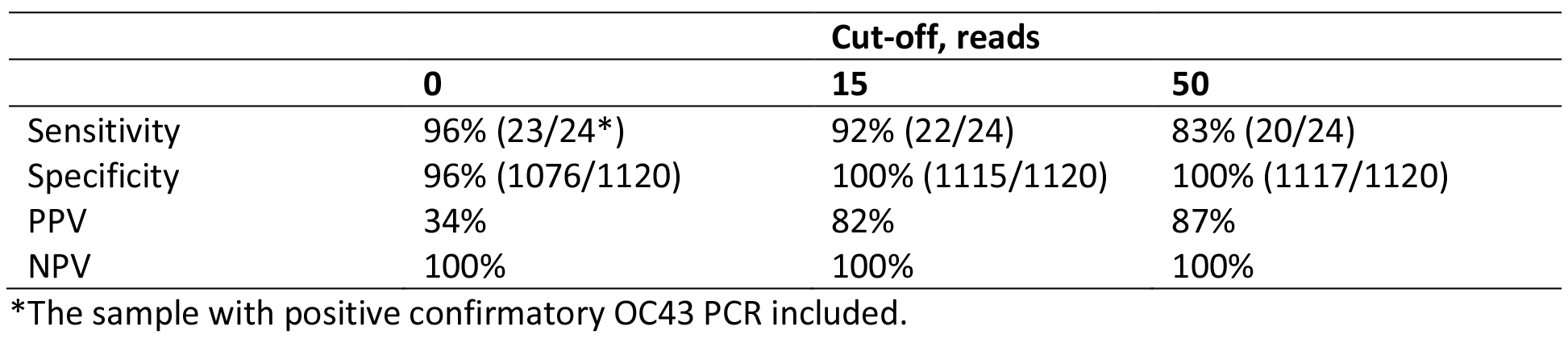
Sensitivity and specificity of mNGS for PCR target viruses. PPV: positive predictive value, NPV, Negative predictive value.

|  | Cut-off, reads |  |  |
| --- | --- | --- | --- |
|  | 0 | 15 | 50 |
| Sensitivity | 96% (23/24*) | 92% (22/24) | 83% (20/24) |
| Specificity | 96% (1076/1120) | 100% (1115/1120) | 100% (1117/1120) |
| PPV | 34% | 82% | 87% |
| NPV | 100% | 100% | 100% |

\*The sample with positive confirmatory OC43 PCR included.

### Typing

mNGS provides additional typing data, as compared to qPCR. Of the 13 rhinoviruses detected with mNGS, 6 (46.2%) were rhinovirus A, 2 (15.4%) rhinovirus B and 5 (38.5%) rhinovirus C. The three influenza viruses were assigned to be H3N2 strains by mNGS.

### Semi-quantification by means of mNGS read count

In order to analyse the semi-quantitative quality of the mNGS assay, the number of the sequence reads (log) mapping to qPCR target viruses (species level) as obtained with mNGS were compared to the Cq values of qPCR. A significant negative correlation was found (Figure 2; Pearson correlation coefficient ρ=−0.5, p=0.008).

**Figure 2:**
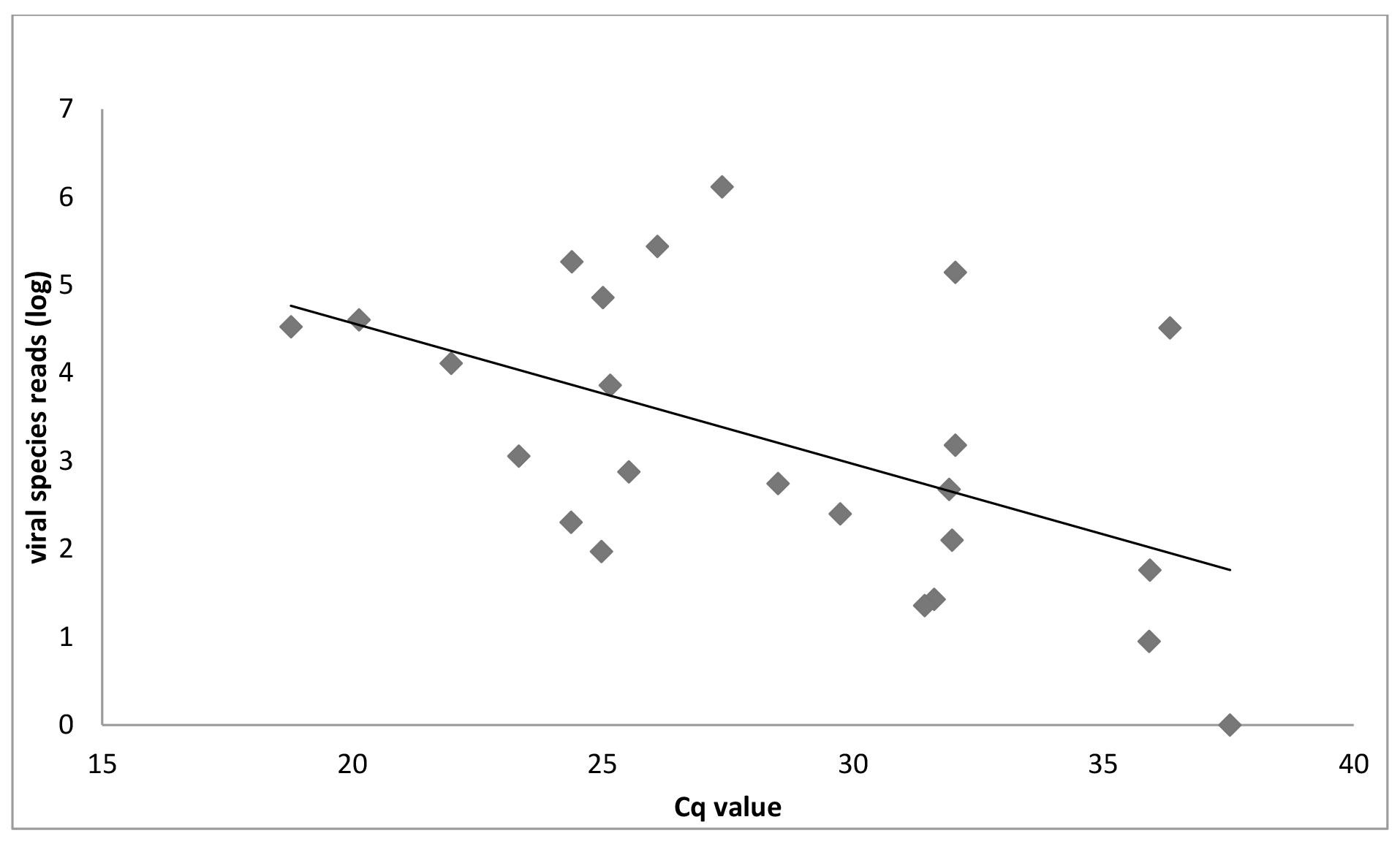
Correlation between mNGS viral species reads (log) and Cq value (ρ=−0.5, p=0.008)

### Clinical parameters and mNGS pathogen read count

The following markers were tested for potential associations with clinical severity of exacerbation (exacerbation severity, self-reported exacerbation severity), length of exacerbation and a decrease/increase in FEVi (control visit compared to baseline): mNGS pathogen positive versus negative exacerbation (qPCR targets), the number of species reads (log) for the different target viruses (species and family level), the number of target virus genus reads (%) of all virus reads. No correlation was found between these markers and the different disease severity parameters (results not shown).

### The respiratory virome

Overall viral families detected by mNGS and abundance of mNGS reads for these families are shown in Figure 3a and b (bacteriophages). Patients with viral pathogens (PCR target viruses) had significantly reduced amounts of bacteriophages when compared to patients without viral pathogen: 17% and 54% bacteriophages respectively (P<0.000, bacteriophage reads vs. all viral reads, excluding EAV control reads). Furthermore, Shannon diversity scores were significantly higher for COPD exacerbations of viral etiology (p<0.000, viral reads in PCR positive versus negative patients, Figure 4).

**Figure 3.**
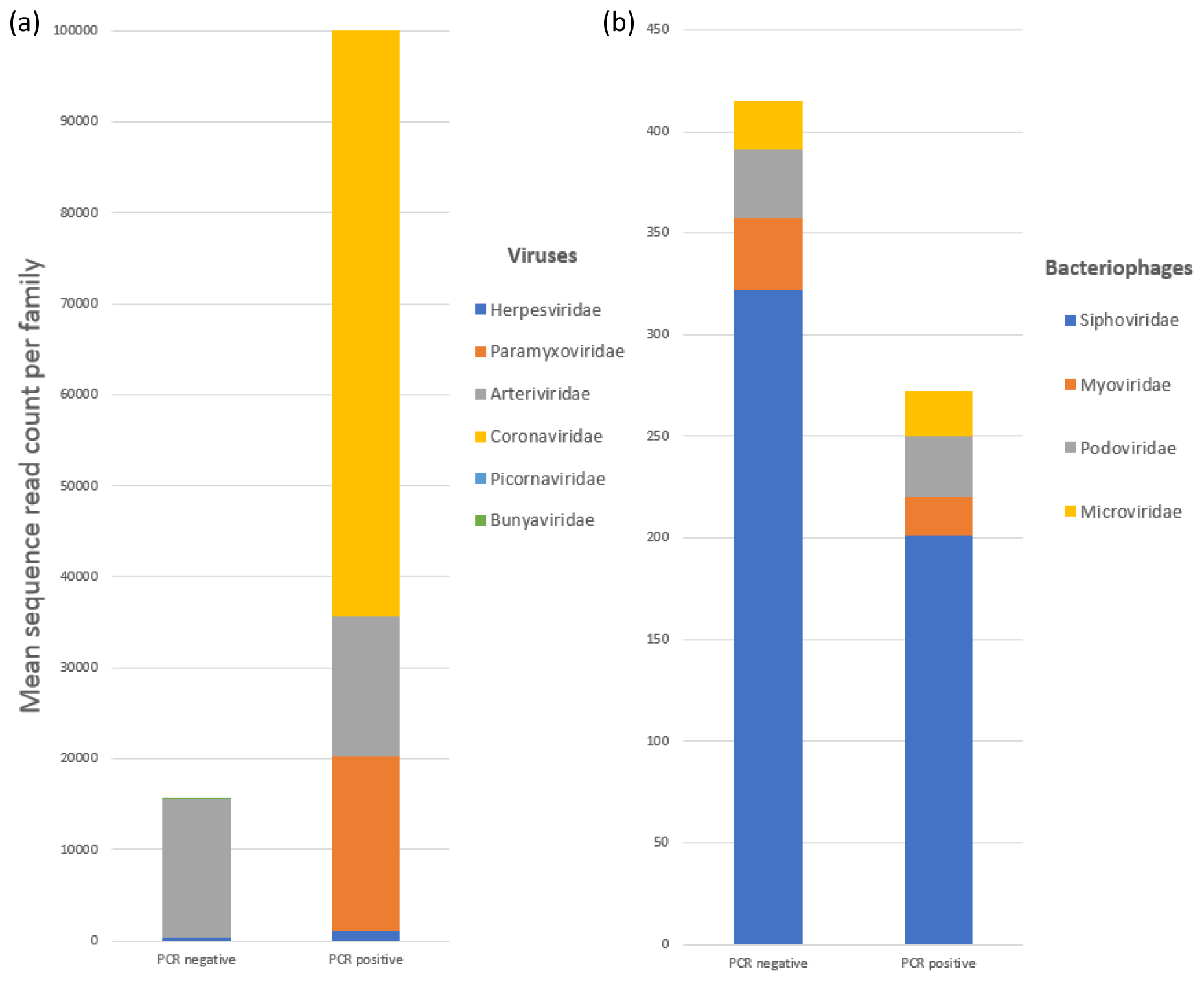
The respiratory virome: abundancy of (a) viral families (bacteriophages excluded) and (b) bacteriophages. Mean sequence read counts per family. Arteriviridae and Herpesviridae include internal control reads (EAV and PhHV-1 respectively). Patients with viral PCR pathogens had lower amounts of bacteriophages (p<0.000).

**Figure 4.**
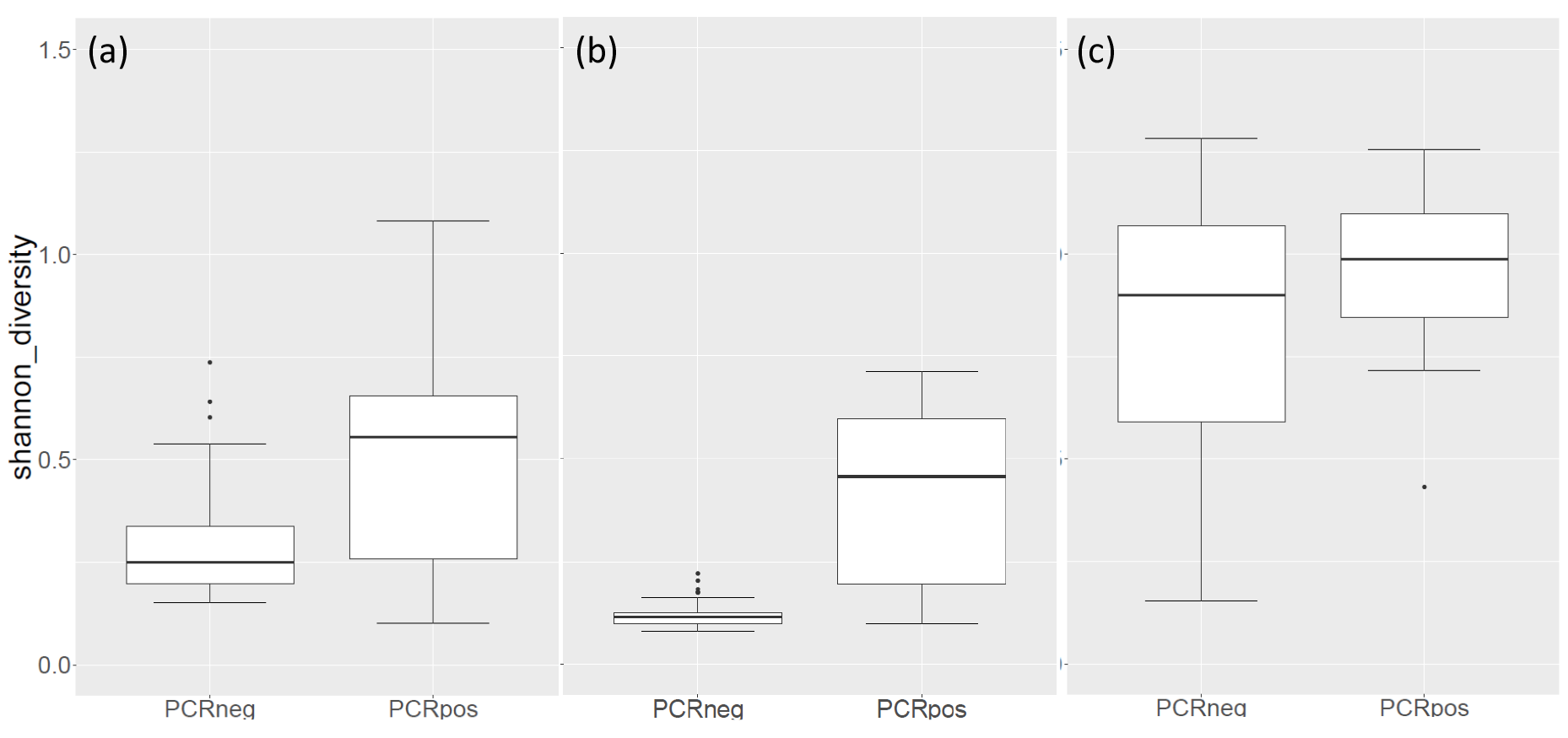
Shannon diversity scores for: (a) viruses, (b) viral PCR targets, (c) bacteriophages. COPD exacerbations of viral etiology had significant lower diversity (b). Boxes span IQR.

No significant association was found between the diversity scores, nor the percentage of bacteriophages, and the following parameters: disease severity, length of exacerbation, number of exacerbations during the study period, difference in FEVi, GOLD stage, smoking, CRP level, and the virus species (results not known).

## Discussion

In this study, the respiratory virome in patients with COPD exacerbations was analysed with both mNGS and qPCR, and combined with clinical data. The incidence of viral pathogens was 26% with both mNGS and qPCR: mNGS failed to detect one Rhinovirus with low load (Cq 38) and PCR failed once due to one of the limitations of PCR, *i.e.* inhibition of amplification. One additional viral pathogen was detected: herpes simplex virus 1, found by others to be associated with COPD^25^.

The incidence of viral pathogens was comparable to that in previous publications (22-64% ^3,5,6^). The viral pathogen with the highest incidence was rhinovirus, followed by influenza, coronaviruses and para-influenza viruses. Interestingly, subtyping data was readily available by mNGS, accentuating the high resolution of mNGS, with rhinovirus RV species A and C being most frequent, followed by RV-B. RV-C was first identified in 2006 and associated with high symptom burdens in children and asthmatics^26,27^. Recently, an asthma-related cadherin-related family member 3 (CDHR3) gene variant^28^ was associated with greater RV-C receptor display on pulmonary cell surfaces of children and adults, and associated with higher susceptibility to severe virus-triggered asthma episodes^29,30^. In line, Romero-Espinoza et al detected predominantly RV-C in children with acute asthma exacerbations by mNGS^31^. The significance of RV-C infection in the adult population is less well studied. Although RV-C has been previously associated with exacerbations of COPD^32,33^, to our knowledge, to date, CDHR3 polymorphisms have not yet found to be associated with COPD.

Furthermore, the complete respiratory virome showed a high phage abundance that could be linked to the absence of viral pathogens. Lower phage abundance may be the result of viral expansion. Hypothetically, a healthy virome size and diversity fits a certain size and diversity of bacteriophages, while during viral infection, pathogens predominate the virome. Alternatively, others have hypothesized that viral and microbial diversity may play a role in infection susceptibility and the development of acute and chronic respiratory diseases^31^. Others have found a higher phage abundance in patients with severe COPD when compared with moderate COPD and healthy controls, in line with the hypothesis of a state of dysbiosis that increases with disease progression^25^. In COPD patients, viral infections have been suggested to trigger bacterial overgrowth and infections^34,35^, demonstrating the significance of viral-bacterial interactions. Moreover, hypothetically, bacteriophages play a role in the horizontal gene transfer of bacterial virulence factors. Study of the lower airways by means of e.g. protected brushes during bronchoscopy are needed for further analysis of bacterial and viral (sub)populations.

The sensitivity, specificity and positive and negative predictive values of mNGS were high: 92%, 100%, 82% and 100%, respectively, when encountering a cut-off of >15 sequence reads, with a detection limit of approximately Cq 38. The high negative predictive value implicates the power of mNGS to exclude any viral infectious cause in one test. The potential to exclude any infectious cause, both viral and bacterial, would have significant consequences for starting and/or continuation of antimicrobial or, at the other end of the spectrum, immune-modulating treatment. The viral species sequence read count might give an indication of the viral burden and the clinical relevancy of the detected virus. Although in our dataset we could not find any correlation with disease severity, several paediatric studies demonstrated a correlation between virus load and disease severity in respiratory infections ^36–39^. Further analysis with a larger number of infected patients and/or a different spectrum of exacerbation severity will be needed to demonstrate or exclude such an association in COPD patients.

Though mNGS renders the possibility to detect all viruses in direct respiratory material, this revolutionary method is not yet used as routine accredited diagnostic procedure for pathogen detection. Before mNGS can be implemented as a routine diagnostics, the optimal protocol must be defined and analysis and interpretation of the metagenomic data must be standardized, followed by external quality assessment. This study demonstrates good performance of our mNGS protocol, in line with other studies ^40,41^ and seems to overcome some of the current thresholds for implementation in clinical diagnostics.

## Conclusions

The mNGS protocol used was highly sensitive and specific for semi-quantitative detection of respiratory viruses. Excellent negative predictive value implicates the power of mNGS to exclude any infectious cause in one test, with consequences for clinical decision making. Reduced abundance of bacteriophages in COPD patients with viral pathogens implicates skewing of the virome, and speculatively the bacterial population, during infection.

## Acknowledgements

We thank our GENERADE project partners Floyd Wittink, Wouter Suring (Hogeschool Leiden), Danny Duijsings (BaseClear) and Christiaan Henkel (Leiden University). We also like to thank Tom Vreeswijk, Lopje Höcker and Mario van Bussel (KML, LUMC) for help with the pre-sequencing experiments and Jeroen Laros (Human Genetics, LUMC) for help with the bioinformatics.

## Disclosures

### Competing interests

none declared.

No part of the manuscript has been previously published.

## Author’s contributions

Initial patient inclusion and sample selection was performed by TME. ALR, SB, ECJC, TME, PB, MA, PSH, ACMK and JJCV participated in the study design. MA examined all patients. ALR performed qPCR testing, ALR, SB, ECC and IS analysed and interpreted the mNGS data. Analyses and interpretation of combined data was performed by ALR, SB, ECC, ECJC, TME, PSH, ACMK and JJCV. ALR wrote the first version of the manuscript. SB,ECC, ECJC, TME, PSH, ACMK and JJCV contributed and revised the manuscript. All authors read and approved the final manuscript.

